# Conditional *Lpar1* gene targeting identifies cell types mediating neuropathic pain

**DOI:** 10.1101/2020.02.02.931212

**Authors:** Richard R. Rivera, Mu-En Lin, Emily C. Bornhop, Jerold Chun

**Author notes:** These authors contributed equally to this work (Richard R. Rivera, Mu-En Lin). RevMAb Biosciences, 830 Dubuque Ave, South San Francisco, CA, USA 94080. To whom correspondence should be addressed: Jerold Chun: Sanford Burnham Prebys Medical Discovery Institute, La Jolla, CA 90237.

## Abstract

LPA_1_ is one of the six known receptors (LPA_1-6_) for lysophosphatidic acid (LPA). Constitutive *Lpar1* null mutant mice have been instrumental in identifying roles for LPA-LPA_1_ signaling in neurobiological processes, brain development, and behavior as well as modelling human neurological diseases like neuropathic pain. Constitutive *Lpar1* null mutant mice are protected from partial sciatic nerve ligation (PSNL)-induced neuropathic pain, however *Lpar1* expressing cell types that are functionally responsible for mediating this protective effect are unknown. Here we report generation of a *Lpar1^flox/flox^* conditional null mutant mouse that allows cre-mediated conditional deletion combined with its use in a PSNL pain model. *Lpar1^flox/flox^* mice were crossed with *cre* transgenic lines driven by neural gene promoters for *nestin* (all neural cells), *synapsin* (neurons), or *P0* (Schwann cells). *CD11b*-*cre* transgenic mice were also used to delete *Lpar1* in microglia. PSNL-initiated pain responses were reduced following cre-mediated *Lpar1* deletion with all 3 neural promoters but not the microglial promoter, supporting involvement of Schwann cells and central and/or peripheral neurons in mediating pain. Interestingly, rescue responses that were due to conditional deletion were non-identical, implicating distinct roles for *Lpar1*-expressing cell types. Our results with a new *Lpar1* conditional mouse mutant expand an understanding of LPA_1_ signaling in the PSNL model of neuropathic pain.

---

Neuropathic pain is produced by nerve lesions or neurological conditions such as multiple sclerosis, diabetes, and cancer affecting an estimated 10% of the general population (1). Treatment options for individuals affected by neuropathic pain are limited and ineffective, often leading to a worsened condition and disability. Initiation and propagation of pain signaling occurs through afferent nerve fibers that relay peripheral signals through dorsal root ganglia (DRG) to signal centrally via the central nervous system (CNS) spinal cord dorsal horn and brain (reviewed in (2–4)). Neuropathic pain involves central sensitization, a process that results in allodynia (painful response to normally innocuous stimuli) and hyperalgesia (increased pain sensation to noxious stimuli) (5).

One identified modulator of neuropathic pain is the bioactive lipid lysophosphatidic acid (LPA). LPA normally signals through six known G protein-coupled receptors, *Lpar1-6* (6), which are involved in myriad biological and pathological processes affecting most of the physiological systems in the body, including the nervous system (6–13). LPA_1_ is also expressed in the peripheral nervous system (PNS) and CNS. Schwann cells represent one of the LPA_1_ expressing cell types that may be involved in the induction of neuropathic pain. LPA signaling through this receptor influences Schwann cell morphology, migration, and survival (14,15). *In-vivo*, sciatic nerves of *Lpar1* deficient mice show abnormalities including an increased number of apoptotic Schwann cells, reduced myelin thickness, and a proportionately lower number of small nerve fiber interacting Schwann cells (14,16). Neurons can also be affected through regulation of neuronal cell morphology, motility, growth cone collapse, calcium signaling, and proliferation (16–23). Mice deficient for this receptor display alterations in cortical development and neurogenesis and show behavioral abnormalities (22–24).

A role for LPA in pain sensation was first identified through intrathecal (i.t.) injection of LPA, where mice that received a single (i.t.) injection of LPA developed thermal hyperalgesia and mechanical allodynia (25). LPA-induced neuropathic pain was accompanied by other sequelae including demyelination in the dorsal root and increased expression of the pain associated markers, protein kinase C γ (PKCγ) in the spinal cord dorsal horn, and voltage-gated calcium channel Caα2δ1 in the DRG (25). Interestingly, i.t. injection of LPA also induced *de novo* production of LPA in the dorsal horn and dorsal root, implicating a feed-forward role in pain generation (26). *De novo* LPA production was also observed in the dorsal horn and dorsal root following PSNL (27–29). Wildtype mice subject to PSNL displayed pain behaviors similar to those of mice that received LPA i.t. and showed similar demyelination as well as upregulation of PKCγ and Caα2δ1 (25).

LPA’s effects in PSNL were shown to be receptor-dependent through the use of constitutive null receptor mutants. *Lpar1* null mutant mice were protected from PSNL and i.t. LPA injection induced mechanical allodynia, and did not show accompanying increased levels of PKCγ and Caα2δ1 (25). *Lpar5* null mutant mice were also protected from PSNL induced neuropathic pain, albeit through CNS mechanisms distinct from those of *Lpar1* null mutants (30).

While *Lpar1* null mutant mice are protected from PSNL-induced neuropathic pain, the cell types responsible for mediating this protection remain unclear. To address this issue, we generated an *Lpar1* conditional null mutant mouse and targeted deletion of *Lpar1* in all neural lineages, peripheral and CNS neurons, Schwann cells, and microglia/macrophages to identify the cell types responsible for mediating *Lpar1*’s protective effect in the PSNL neuropathic pain model.

## Results

### Generation of *Lpar1* conditional null mutant mice

We selected a portion of the *Lpar1* genomic locus for conditional gene targeting in embryonic stem (ES) cells resulting in the creation of a mutant mouse, designated *Lpar1^flox/flox^*, where *Lpar1* exon 3 is selectively deleted in the presence of the cre recombinase (Fig. 1A). The targeting vector contained a loxP site that was introduced into a restriction enzyme site 5’ of exon 3, and a neomycin drug selection cassette flanked by loxP sites in a restriction enzyme site 3’ of exon 3. Following electroporation of the linearized *Lpar1* targeting construct, drug selection, and screening of DNA isolated from selected ES cell clones by Southern blotting and hybridization, several clones with a homologously recombined *Lpar1* allele were identified (Fig. 1B and 1C). PCR with primers flanking the 5’ loxP site was used to select ES cell clones for blastocyst injection. Mice positive for germline transmission of the recombined allele were then crossed with *nestin-cre* transgenic mice to produce *Lpar1^flox/flox^*-*nestin*-*cre* mice (31). Because *nestin* is expressed in the testis, male *Lpar1^flox/flox^*-*nestin*-*cre* mice were bred to C57BL/6J female mice to produce offspring with germline *cre*-mediated loxP site recombination. Selective deletion of the floxed neomycin cassette and retention of the 5’ loxP site in offspring were identified by PCR (Fig. 2A and 2B). Heterozygous *Lpar1*^*flox*/+^ mice with the correct recombination events were then crossed together to produce wildtype, *Lpar1*^*flox*/+^, and *Lpar1^flox/flox^* mice (Fig. 2C). A high level of embryonic lethality was observed for *Lpar1* constitutive null mutant mice in a C57BL/6J background whereas *Lpar1^flox/flox^* mice in this background strain are healthy and are indistinguishable from wildtype littermates. Wildtype, *Lpar1*^*flox*/+^, and *Lpar1^flox/flox^* mice were identified by PCR (Fig. 2C) and are behaviorally the same.

**Figure 1.**
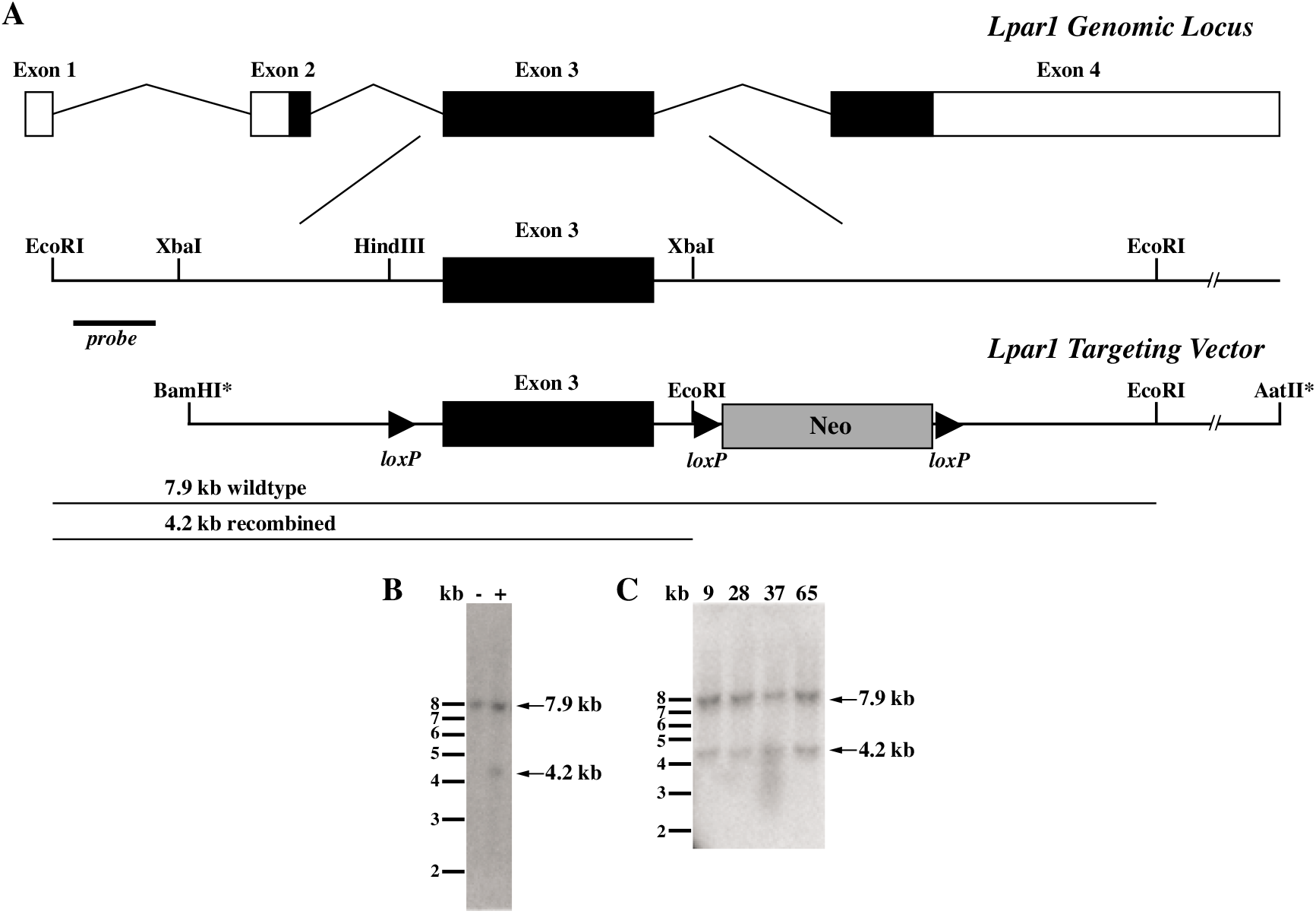
Conditional gene targeting of the *Lpar1* gene locus and identification of ES cells positive for homologous recombination. (A) Schematic of the *Lpar1* genomic locus, the region used for gene targeting, and the *Lpar1* targeting vector. In the targeting vector, loxP sites flank *Lpar1* exon 3 and the neomycin cassette used for ES cell drug resistance selection screening, an introduced EcoRI site allows for identification of homologous recombination events with the indicated external probe. Asterisks represent artificial restriction enzyme sites used in the construction of the targeting vector. (B) Southern blot of EcoRI digested ES cell DNA hybridized with the radiolabeled probe shown in (A) identifies an ES cell clone positive (+) for homologous recombination, as indicated by the presence of a 4.2 kb band. An ES cell clone with an incorrect recombination event (-) is shown for comparison and shows only the wildtype 7.9 kb *Lpar1* band. (C) Four identified ES cell clones (9,28,37, and 65) were grown and homologous recombination was reconfirmed by Southern blotting. These clones were chosen for loxP site retention screening, clones 37 and 65 were used for used for blastocyst injections.

**Figure 2.**
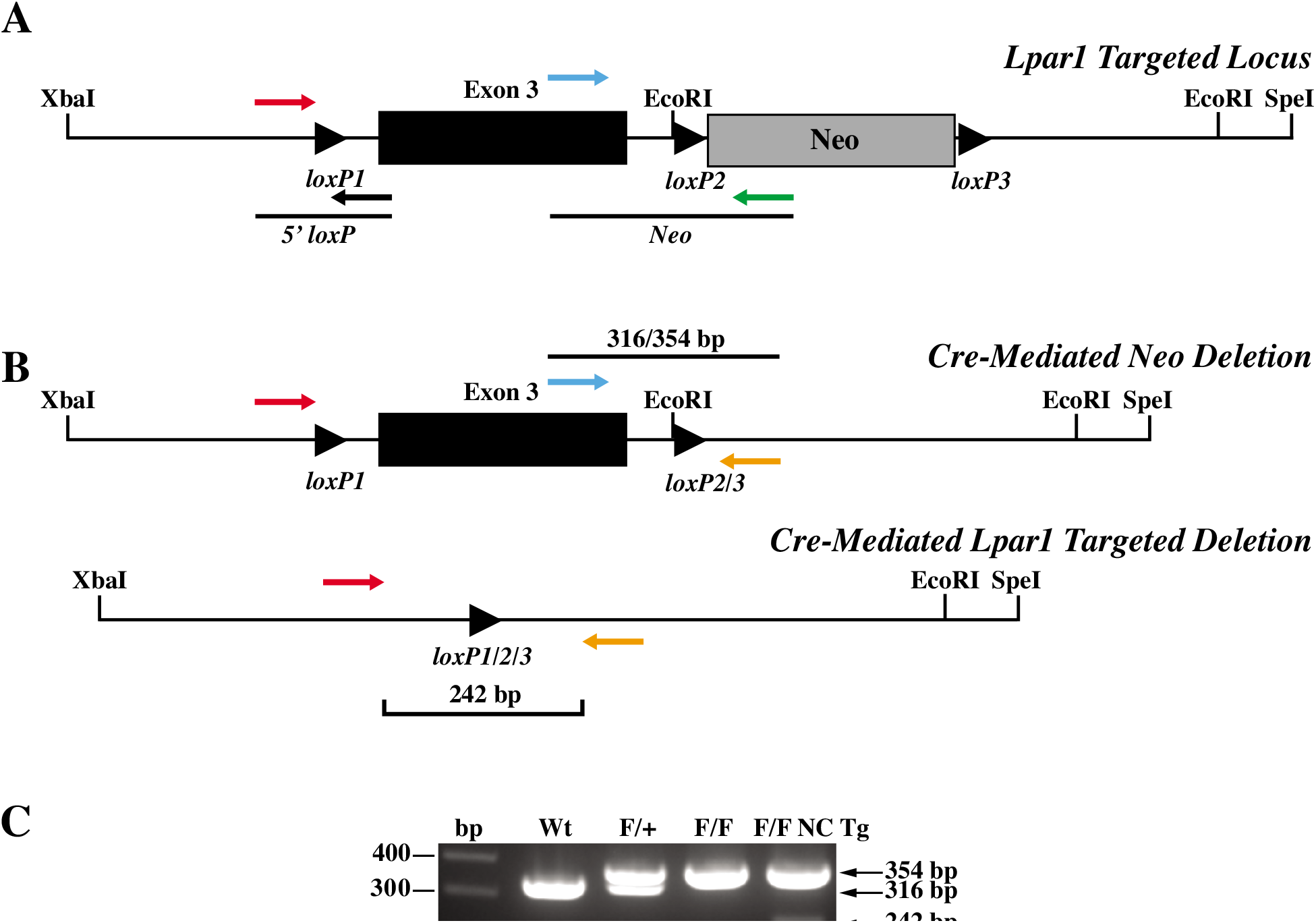
Cre-mediated deletion in mice with a recombined *Lpar1* allele. (A) Schematic showing PCR primer pairs used to screen for cre-mediated deletion of the floxed neomycin cassette. The primers shown assay for the presence of the 5’ loxP site and the presence or absence of the neomycin cassette. (B) Diagrams showing the finished *Lpar1* targeted allele produced through *in vivo* cre-mediated deletion of the neomycin cassette (top) and cre-mediated targeted deletion of floxed *Lpar1* exon 3 (bottom). The three-primer combination used for PCR genotyping is indicated. (C) PCR genotyping of tail DNA from wildtype (Wt), *Lpar1*^*flox*/+^(F/+), *Lpar1^flox/flox^*(F/F), and *Lpar1^flox/flox^*-*nestin-cre* transgenic (F/F NC Tg) mice. Primers shown in (B) were used for PCR. Wildtype *Lpar1* produced bands of 316 bp, while floxed alleles produced bands 354 bp. The presence of a 242 bp band in the *Lpar1^flox/flox^*-*nestin-cre* sample is indicative of cre-mediated deletion in neural tissue present in the mouse tail.

### Cre-mediated *Lpar1* targeted deletion

To delete *Lpar1* in all neural cell types, neurons, Schwann cells, and myeloid lineage cells, *Lpar1^floxflox^* mice were crossed to *nestin, synapsin, P0*, and *CD11b*-*cre* transgenic mice, respectively (31–34). To confirm that *Lpar1* was deleted in the presence of cre, genomic DNA was isolated from DRG of *Lpar1^flox/flox^* and *Lpar1^flox/flox^*-*nestin*-*cre* mice and PCR was used to verify genomic recombination of the *Lpar1* genomic locus to produce a null allele (Fig. 3A). DRG contain both neural and non-neural cells, with conditional deletion limited to neural cells, thus producing a recombined (neural) and unrecombined (non-neural) signal in conditional mutants. As expected, PCR products indicative of both an unrecombined and recombined *Lpar1^flox/flox^* allele can be amplified from genomic DNA isolated from *Lpar1^flox/flox^*-*nestin-cre* DRG, while only an unrecombined product can be produced from the DRG of control *Lpar1^flox/flox^* mice (Fig. 3A). In agreement with genomic deletion of *Lpar1*, RT-PCR showed *Lpar1* mRNA transcripts are absent in *Lpar1^flox/flox^*-*nestin-cre* DRG (Fig. 3B). Following Schwann cell specific *P0 cre* crossing, PCR analyses of sciatic nerve showed deletion of *Lpar1* (Fig. 3C), compared to wildtype. Neuronal deletion was confirmed in cerebral cortex (Ctx) of *Lpar1^flox/flox^*-*synapsin-cre* mice (Fig. 3D). Immunofluorescent labeling of peripheral myelinated axons for MBP (myelin in red) and satellite glia expressing LPA_1_ (green) in wildtype DRG (Fig. 3E) was not observed in *Lpar1^flox/flox^*-*nestin-cre* mice (Fig. 3F). These data demonstrate conditional deletion of *Lpar1* in the presence of targeted cre recombinase expression.

**Figure 3.**
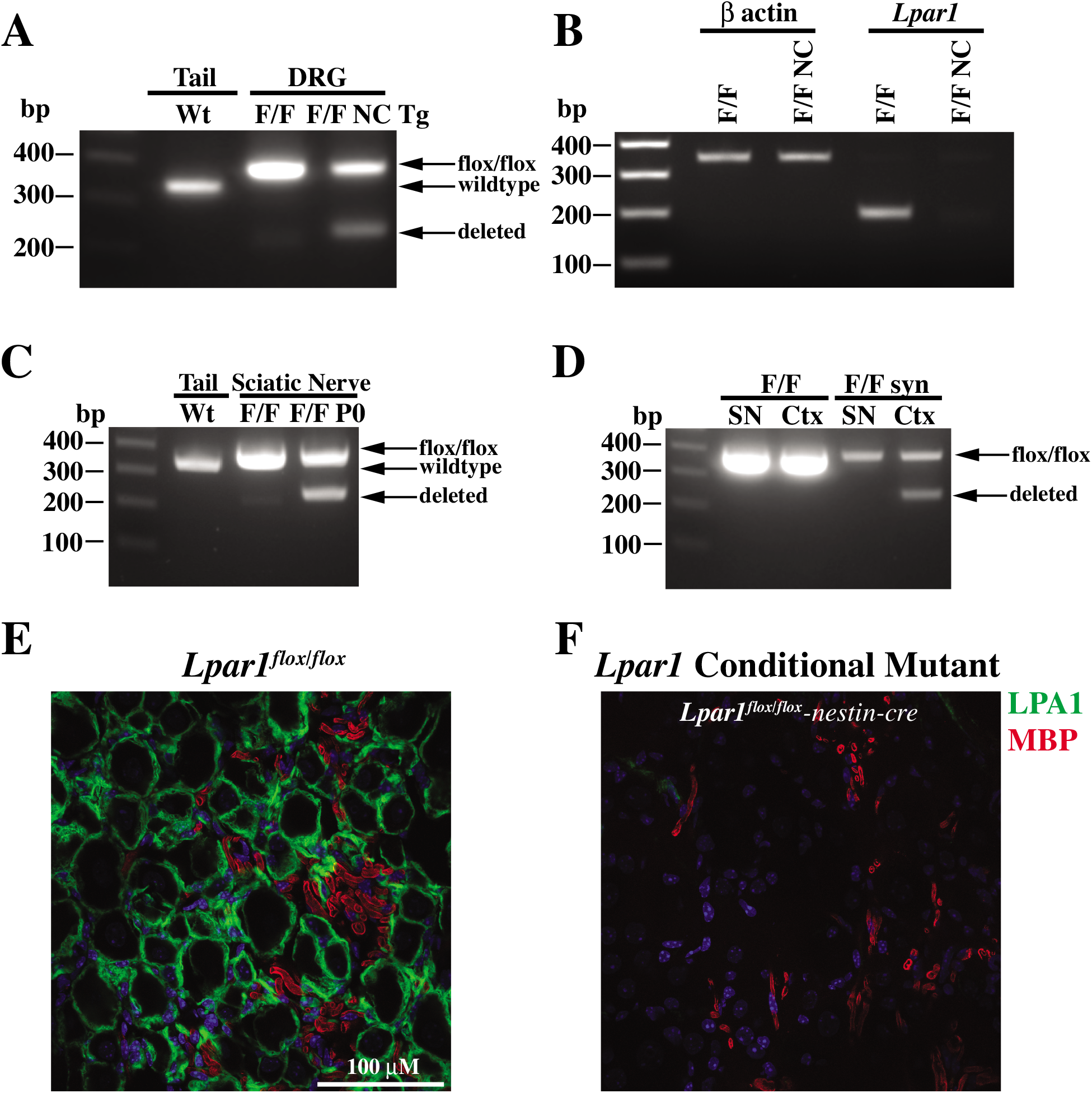
Functional deletion of *Lpar1* is cre-dependent. (A) PCR products of DNA isolated from *Lpar1^flox/flox^* (F/F) and *Lpar1^flox/flox^*-*nestin-cre* transgenic (F/F NC Tg) DRG shows genomic deletion of *Lpar1* exon 3, DNA from the tail of a wildtype mouse is shown for comparison. (B) qPCR products of cDNA prepared from *Lpar1^flox/flox^* (F/F) and *Lpar1^flox/flox^*-*nestin-cre* transgenic (F/F NC Tg) DRG shows that *Lpar1* transcripts are lost in neural tissues. (C) PCR of genomic DNA isolated from the tail of a wildtype mouse (Wt) and the sciatic nerve of *Lpar1^flox/flox^* (F/F) and *Lpar1^flox/flox^*-*P0-cre* (F/F P0) mice. (D) PCR amplification of genomic DNA isolated from the sciatic nerve (SN) and cortex (Ctx) of *Lpar1^flox/flox^* and *Lpar1^flox/flox^*-*synapsin-cre* mice. The PCR primers used for amplification of wildtype (316 bp), *Lpar1* floxed alleles (354 bp), and *Lpar1* deleted products (242 bp), are identical to those used in Fig. 2B. The 100, 200, 300, and 400 bp bands of the 1kb plus DNA ladder are indicated for reference. (E, F) Immunofluorescent labeling of peripheral myelinated axons identify wildtype LPA_1_ immunolabeling in *Lpar1^flox/flox^* mice (E) and its absence (F) in *Lpar1^flox/flox^-nestin-cre* transgenic mice. LPA_1_ labeling is in green and MBP (myelin) in red for individual samples. Scale bar = 100 μM

### *Lpar1* expressing neural cell types contribute to PSNL-induced neuropathic pain phenotypes

To determine which *Lpar1* expressing neural cell types mediate PSNL-induced neuropathic pain protection, paw withdrawal threshold responses following cre recombination for *Lpar1*^flox/flox^-*nestin*, *Lpar1*^flox/flox^-*synapsin*, *Lpar1*^flox/flox^-*P0* and *Lpar1*^flox/flox^-*CD11b*-*cre* was assessed. No pain rescue was observed with *Lpar1*^flox/flox^-*CD11b* compared to wildtype or to *Lpar1^flox/flox^* transgenic mice (data not shown). All other genotypes rescued the pain phenotype compared to controls (Fig. 4A-D). *Lpar1^flox/flox^*-*nestin-cre* conditional mutant mice challenged with PSNL had similar paw withdrawal threshold responses compared to previously defined *Lpar1* constitutive null mutant mice (25) (Fig. 4A). By contrast, *Lpar1^flox/flox^*-*P0-cre* mice initially responded like control mice at early time points (days 3 and 6), but then showed sustained protection at later time points (day 9 through day 21) (Fig. 4B and 4D). *Lpar1^flox/flox^*-*synapsin-cre* mice were initially refractory to PSNL-induced neuropathic pain (Fig. 4C) but lost protection over time (day 12 through day 21) (Fig. 4D). It is notable that the combined protection of *P0* and *synapsin*-*cre* recombination approximated the protection produced by *nestin-cre* recombination (Fig. 4A), implicating an additive rescue effect produced by both Schwann cells and neuronal LPA_1_ activation in PSNL-initiated pain.

**Figure 4.**
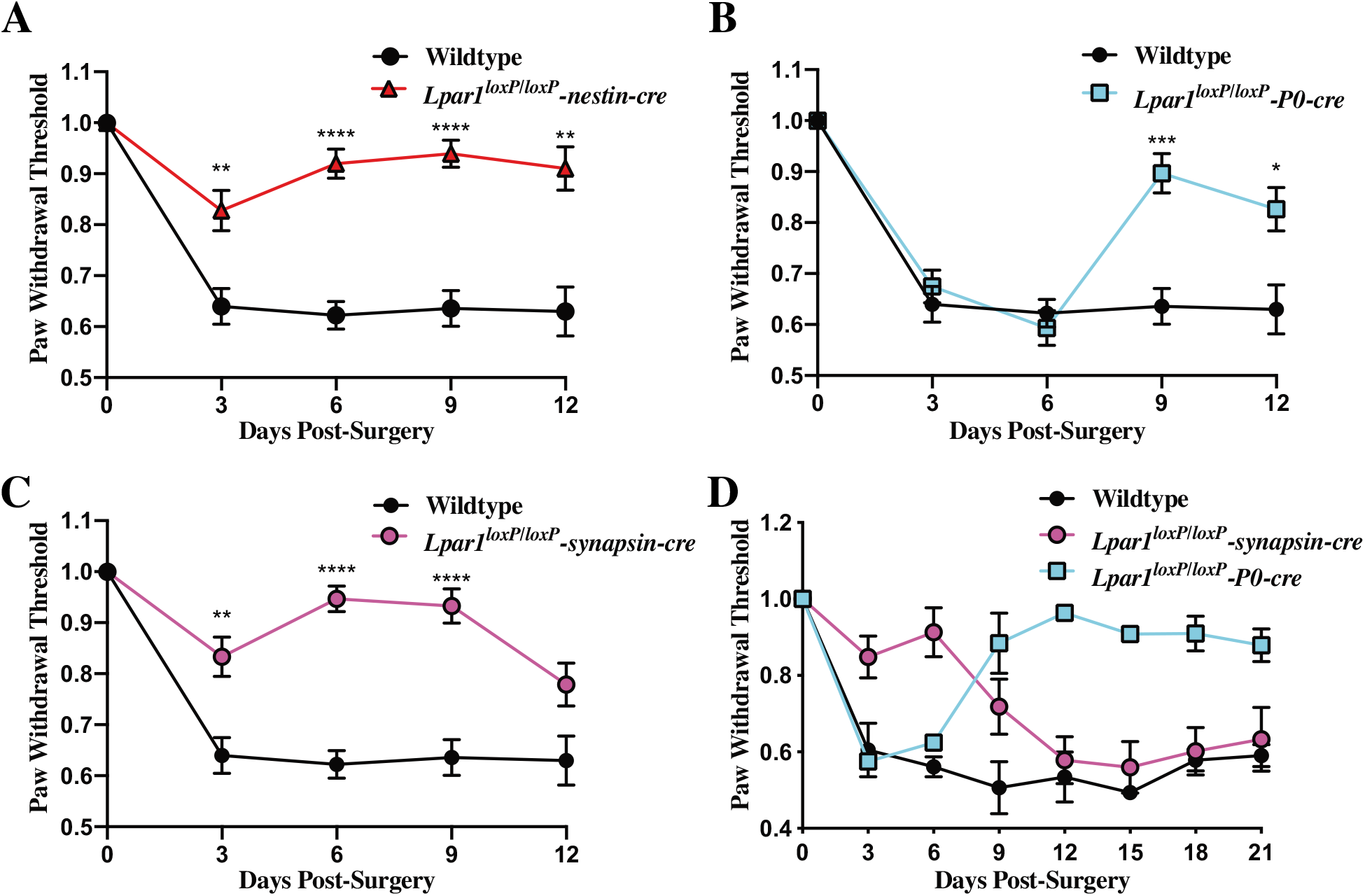
Deletion of *Lpar1* in neuronal lineages protects against PSNL induced neuropathic pain. (A) Targeted *nestin-cre*-mediated deletion of *Lpar1* in all neural lineages protects against neuropathic pain in the PSNL mouse model. (B) Schwann cell specific deletion of *Lpar1* through a *P0-cre* transgene protects mice from PSNL at later but not earlier time points. (C) Specific deletion of *Lpar1* in neurons protects mice from PSNL induced neuropathic pain at early time points but not at later time points. (D) Schwann cell specific deletion of *Lpar1* occurs at later time points and is long-lasting. The plotted data is the average paw withdrawal threshold time observed for *Lpar1* conditional null mutants normalized to *Lpar1^flox/flox^* control animal responses +/- SEM. For (A, B, and C), N=10 *Lpar1^flox/flox^*, N=10 *Lpar1^flox/flox^*-*nestin-cre*, N=9 *Lpar1^flox/flox^*-P0 cre, and N=8 *Lpar1^flox/flox^*-*synapsin-cre* animals. In (D), N=2 for all genotypes used. Statistical analysis was performed using a two-way Anova, followed by a Sidak’s multiple comparisons test, differences were considered significant when P≤0.05 (*=P≤0.05, **≤.001, ***≤0.001, ****P≤0.0001).

## Discussion

*Lpar1* conditional null mutant mice were generated and shown to undergo cre-mediated recombination, enabling identification of *Lpar1*-expressing neurons and Schwann cells as functionally important for the PSNL phenotype. In the absence of cre, *Lpar1^flox/flox^* mice developed a pain phenotype comparable to *Lpar1* constitutive null mutant mice (25), demonstrating that this new *floxed* mutant gene functions normally in PSNL before *cre* crossing. *Lpar1^flox/flox^*-*nestin-cre* mice with a pan-neural lineage deletion of *Lpar1* are protected from PSNL-induced neuropathic pain, supporting neural LPA_1_ signaling as important despite *Lpar1’s* ubiquitous tissue expression. By comparison, *P0 and synapsin-cre* recombination produced only partial rescue with complementary temporal phases of protection that appeared additive to account for the degree of rescue by *nestin-cre* recombination.

The actions of LPA_1_ in Schwann cells affecting PSNL phenotypes have not, to our knowledge, been previously reported, and the observed phenotype was unexpected with regard to the clear and differential time-dependence of the effect. Explanations for these temporal changes in pain protection may be due to differences in *de novo* synthesis of LPA and the varied activation states documented for LPA_1_ (8,11,35–38) that may occur in neurons and Schwann cells. Such LPA signaling effects could be altered by receptor removal to produce the time-course differences observed for PSNL-initiated pain rescue. Long-lasting protection from neuropathic pain at later time points may also reflect changes in nerve myelination that may interfere with the transmission of pain stimuli as previously suggested (14,25). Nerve fibers in *Lpar1^flox/flox^*-*P0-cre* mice may already be abnormally myelinated, and nerve injury induced demyelination may alter normal pain signal transmission. However, we note that the nerve fibers that respond to noxious stimuli are lightly myelinated Aβ fibers and unmyelinated C-fibers (2,3), requiring a more complex scenario that might involve central pain consolidation through myelinated fibers.

Effects of *Lpar1* deletion from neurons in *Lpar1^flox/flox^*-*synapsin-cre* mice showed early protection in PSNL, contrasting with later protection of Schwann cell receptor deletion, while supporting the involvement of neurons in LPA_1_-mediated PSNL-induced pain. *Synapsin-cre* deletion is effective in deleting genes from CNS neurons (39) but less effective in peripheral (DRG) neurons (33,40), implicating central neuronal mechanisms. A possible explanation for rescue at early timepoints could involve a lack of *de novo* LPA synthesis from *Lpar1* deficient neurons. LPA can be released by neurons following nerve transection and neurons can synthesize LPA *de novo* through an LPA receptor dependent feed-forward mechanism (as evidenced by LPA_3_) (15,26,41). *De novo* LPA synthesis from other cell types following PSNL may result in LPA accumulation to drive neuropathic pain at later time points, particularly through activation of Schwann cell receptors in the neuron-specific mutants. Alternatively, PSNL may cause damage and vascular leakage that exposes peripheral nerves to LPA, activating cognate receptors to produce aberrant pain signaling (9,42–44).

Other LPA receptor subtypes can contribute in distinct ways to neuropathic pain based on analyses of different LPA receptor-null mutants (13,16,25,30,37,45,46). Similar to *Lpar1* null mutant mice, *Lpar5* null mutant mice are protected from PNSL-induced neuropathic pain and also show decreased sensitivity to acute pain stimuli and faster recovery responses when challenged in an inflammatory pain model (30,47). Additionally, deletion of *Lpar3* in mice prevents i.t. LPA-induced *de novo* production of LPA in the dorsal horn and dorsal root and also prevents LPA-induced allodynia and hyperalgesia (26), suggesting an *Lpar3* mediated feed-forward mechanism for LPA in neuropathic pain initiation. Prevention of LPA *de novo* synthesis and neuropathic pain in the i.t. LPA and PSNL neuropathic pain models using minocycline combined with *Lpar3* expression in microglia indicate that this feed-forward mechanism is likely mediated by microglia (48,49). In the present study, we did not observe a rescue effect of *Lpar1* loss from microglia, suggesting that maintained *Lpar3* could sustain PSNL-initiated pain. The actual cell types involved in the functions of these and other LPA receptor subtypes remain to be determined but is experimentally tractable through generation of conditional mutants.

The generated *Lpar1* conditional mutant mice will be useful in identifying cell types involved directly with LPA_1_ signaling in neuropathic pain as well as many other conditions and disease models (6–12,37,45,50,51). The tractability of LPA_1_ as a member of the lysophospholipid receptor family supports its potential as a druggable GPCR target (8,10,35,36) for the development of improved therapies targeting specific *Lpar1* expressing cell types.

## Experimental procedures

### Mice

All procedures performed on animals were IACUC approved and performed in accordance with the regulations of The Scripps Research Institute (TSRI) Department of Animal Resources and the Sanford Burnham Prebys Medical Discovery Institute animal care and use committees. Mice used in this study were *nestin-cre* (Jackson Laboratory Stock Number 003771), *P0-cre* (Jackson Laboratory Stock Number 017927), *synapsin-cre* (Jackson Laboratory Stock Number 003966), and *CD11b-cre* (obtained from Don Cleveland) transgenic lines.

### Synthesis of the *Lpar1* conditional gene targeting vector

Creation of the *Lpar1* conditional gene targeting vector was accomplished by PCR amplification of mouse *Lpar1* genomic fragments using a bacterial artificial chromosome (BAC RP23-149020 Children’s Hospital Oakland Research Institute (CHORI)) containing the *Lpar1* genomic locus as a template. PCR amplification was performed using *Pfx50* DNA polymerase (Invitrogen) and amplified genomic fragments were assembled into pBluescript II. During the process of assembly, a loxP site was inserted into a HindIII site 5’ of *Lpar1* exon 3 and a neomycin cassette under the control of the phosphoglycerate kinase promoter (PGK-neo) flanked by loxP sites was inserted directionally (all loxP sites in the same orientation) into an XbaI site 3’ of *Lpar1* exon 3 (Fig. 1A). The construct was engineered so that 3.4 and 6.7 kb of *Lpar1* genomic DNA flanked the PGK-neo insertion site. To aid in cloning, BamHI and AatII restriction enzyme sites were added to the distal 5’ and 3’ ends of the *Lpar1* genomic segment chosen for targeting vector design. An EcoRI restriction enzyme site was included in the loxP flanked PGK-neo cassette to identify ES cell clones containing an allele that recombined homologously with the targeting vector.

### Production of *Lpar1^flox/flox^* and *Lpar1^flox/flox^*-cell type specific null mutant mice

To create the *Lpar1^flox/flox^* mice, 1 x 10^7^ R1 ES cells were mixed with 50 μg of linearized *Lpar1* targeting vector in a 0.4 cm electroporation cuvette and the cells were pulsed with a Bio-Rad Gene Pulser II (200 mVolts x 800 μF capacitance). The electroporated ES cells were plated on mitotically inactive mouse feeder cells and allowed to recover for 24 hrs at 37°C; 24 hrs after electroporation and plating, 150 μg/ml Geneticin (Invitrogen) was added to the ES cell medium and the cells were placed under selection for 7 days. ES cell clones were then isolated and grown individually for subsequent DNA isolation and screening for homologous recombination events by Southern blotting and hybridization with an *Lpar1* DNA probe containing sequence external to that of the 5’ end of the *Lpar1* targeting vector. Clones with homologous recombination events were then screened for retention of the loxP site 5’ to *Lpar1* exon 3 with the following primers: 5’ loxP Forward 5’-gttgggacatggatgctattc-3’ and 5’ loxP Reverse 5’-aatctgttctcatcccacacg-3’. Correctly targeted ES cell clones were then injected into C57BL/6J blastocysts at the TSRI Murine Genetics Core.

To delete the loxP flanked PGK-neo cassette *in-vivo*, gene targeted mice were crossed to *nestin-cre* transgenic mice and resultant males were then bred to C57BL/6J female mice. Male mice were chosen because cre is expressed in the germline of *nestin-cre* male mice. Offspring were then screened by PCR for the presence of the 5’ loxP site with the primers listed above, in the presence or absence of the PGK-neo cassette with primers A1 Exon 3 Forward 5’-agactgtggtcattgtgcttg-3’ and Neo Reverse 5’-tggatgtggaatgtgtgcgag-3’, and for retention of the loxP site 3’ to *Lpar1* exon 3 with primers 3’ loxP Forward 5’-tgcagaattatgagtggacagg-3’ and 3’ loxP Reverse 5’-ggtttagtggtgtgggatcg-3’. Mice that retained the loxP sites 5’ and 3’ to *Lpar1* exon 3 but deleted the PGK-neo cassette were selected for propagation and crossing with *nestin-cre*, *P0-cre*, *synapsin-cre*, and *CD11b-cre* transgenic mice (31–34).

PCR genotyping of the *Lpar1* conditional mutant mice was done with the following primers: 5’ loxP Forward 5’-gttgggacatggatgctattc-3’, 3’ loxP Reverse 5’-ggtttagtggtgtgggatcg-3’, and A1 Exon 3 Forward 5’-agactgtggtcattgtgcttg-3’. PCR amplification of genomic DNA with these primers identified wildtype, *Lpar1^flox^*, and *Lpar1* deleted products of 316, 354, and 242 bp, respectively. *Synapsin-cre*, *CD11b*, *P0-cre*, and *nestin-cre* transgenes were identified by PCR amplification of genomic DNA with a common reverse PCR primer, (Cre Reverse 5’-CAG CAT TGC TGT CAC TTG GTC-3’), and forward primers specific for *synapsin* (SynCreForward 5’-CCCAAGAAGAAGAGGAAGGTG-3’), *CD11b* (CD11b Forward 5’-ACACCTCAGCCTGTCCAGTAG-3’), P0 (MPZ Forward (P0 Cre) 5’-ATT GGT CAC TGG CTC AAG AC-3’), and *nestin* (Nestin Prom: 5’-ACT CCC TTC TCT AGT GCT CCA-3’) yielding products of 350 bp, 1 kb, 525 bp, and 550 bp respectively.

### Southern blotting and DNA hybridization

ES cell clones were screened for homologous recombination by digesting 10 μg of ES cell DNA with EcoRI, running the DNA on a 0.8% 1 x TAE agarose gel, and transferring the digested DNA to Nytran SuPerCharge membrane (GE Healthcare Life Sciences) in 20 x SSPE. Transferred DNA was UV crosslinked to the membrane and hybridized with a ^32^P-labeled (Prime-It II Random Primer Labeling Kit, Agilent) *Lpar1* probe with sequence external to the 5’ end of the targeting vector. The 800 bp probe was produced by PCR from a BAC containing *Lpar1* with the following primers: A1 Ext Forward 5’-actgaggtcacttactcagag-3’ and A1 Ext Reverse 5’-gtctatggctgtggaattcaag-3’. Probe hybridization was carried out overnight at 42°C in a .05 M pH 7.4 phosphate buffer containing 50% formamide, 5 x SSPE, 1 x Denhart’s, 1% SDS, containing .1% denatured 10 mg/ml salmon sperm DNA following a 1 hr pre-hybridization. Blots were washed and visualized using a phosphorimager. The presence of a 4.2 kb recombined band and a 7.9 kb wildtype band was indicative of ES cells with homologous recombination events.

### Partial sciatic nerve ligation and behavioral testing

The partial sciatic nerve ligation (PSNL) procedure was performed as described (30). Adult *Lpar1^flox/flox^* and *Lpar1^flox/flox^*-*cre* transgenic mice in a C57BL/6J background were anesthetized via nosecone delivery of isoflurane and the right limb sciatic nerve exposed and tightly ligated with 10-0 fine sutures. The wound and skin were closed and stitched, and the animals allowed to recover. For behavioral testing, animals were acclimated in cages with wire mesh bottoms for one hour prior to testing in an environmentally controlled testing room. Paw withdrawal threshold (gram (g)) against increasing mechanical stimuli (0-50 g in 20 s) were measured before and following PSNL surgery with tests conducted four separate times with at least a 1 min interval between tests. The average response was normalized to pre-surgery controls +/- SEM.

### Immunohistochemistry

DRG were isolated from the lumbar region of *Lpar1^flox/flox^* control and *Lpar1^flox/flox^*-conditional null-mutant mice. Tissues were embedded in OCT compound and 5 μM sections were cut and immunolabeled with antibodies to mouse LPA_1_ (PA1 10401, Thermo Fisher Scientific), MAG (clone 513 MAB-1567, Chemicon), MBP (ab134018 Abcam), and TUJ1 (MMS-435P, Covance). Secondary antibodies were used against the listed primary antibodies and 60x images were acquired on a Zeiss Axio Imager.D2 microscope.

### Reverse transcription PCR

DRG were isolated from the lumbar region of *Lpar1^flox/flox^* and *Lpar1*^flox/flox^-*nestin-cre* conditional null-mutant mice. DRG were placed in 1 ml of TRIzol Reagent (Thermo Fisher Scientific) and total RNA was isolated according to the manufacturer’s directions. cDNA was synthesized from total RNA using a Bio-Rad iScript cDNA synthesis kit and β-actin and *Lpar1* specific oligonucleotide primer pairs were used to amplify target gene transcripts. Primers used to amplify a 350 bp product from β actin cDNA were M β Actin Forward 5’-tggaatcctgtggcatccatg-3’ and M β Actin Reverse 5’-aaacgcagctcagtaacagtc-3’; primers used to amplify a 194 bp product from *Lpar1* cDNA were M LPA_1_ Forward RT 5’-gacaccatgatgagccttctg-3’ and M LPA_1_ Reverse RT 5’-tcgcggtaggagtagatgatg-3’. An equivalent amount of cDNA from each sample, calibrated to produce equal amounts of β-actin PCR product, was used to amplify the *Lpar1* cDNA.

## Acknowledgments

We thank Dr. Andras Nagy for the R1 ES cells used for gene targeting, Grace Kennedy for histology expertise, Dr. Gwendolyn Kaeser for statistical analysis, and Dr. Gwendolyn Kaeser and Danielle Jones for editorial assistance.

## FOOTNOTES

Funding was provided by the NIMH of the National Institutes of Health under award number R01MH051699 to J.C. and non-Federal funds from a predoctoral fellowship from Amira Pharmaceuticals to M.L. The content is solely the responsibility of the authors and does not necessarily represent the official views of the National Institutes of Health.

The abbreviations used are:

DRG: dorsal root ganglia
CNS: central nervous system
LPA: lysophosphatidic acid
PNS: peripheral nervous system
PNSL: partial sciatic nerve ligation.

